# Dynamin 1-mediated endocytic recycling of glycosylated N-cadherin sustains the plastic mesenchymal state to promote ovarian cancer metastasis

**DOI:** 10.1101/2024.07.16.603672

**Authors:** Yuee Cai, Sally K. Y. To, Zhangyan Guan, Yin Tong, Jiangwen Zhang, Ling Peng, Philip P. C. Ip, Alice S. T. Wong

## Abstract

Epithelial-to-mesenchymal transition (EMT) is a key process that confers metastatic plasticity to ovarian cancer cells, enabling them to disseminate aggressively throughout the peritoneal cavity and contributing to poor clinical outcomes for patients. However, a pharmacologically exploitable driver of EMT in ovarian cancer has yet to be identified. To address this, we utilized a master regulators algorithm to prioritize EMT regulators from a dataset of over 8,000 patient samples, including multidimensional omics data from more than 20 cancer types in TCGA. Further analysis identified dynamin-1 (DNM1), an endocytic regulator, as a novel master regulator of EMT in ovarian cancer. Clinically, DNM1 overexpression was found to be associated with the mesenchymal subtype and advanced/metastatic stages of ovarian carcinomas. Molecular assays revealed that DNM1 upregulates N-cadherin, a hallmark mesenchymal marker, by promoting its endocytosis and recycling, thereby inducing cell polarization and motility. In addition, integration of ATAC-seq and RNA-seq analyses uncovered the repression of beta-1,3-galactosyltransferase (B3GALT1), a glycosyltransferase, in metastatic cells. B3GALT1-mediated glycosylation hindered the recycling of N-cadherin. Functional studies demonstrated that depletion of DNM1 or pharmacological inhibition of endocytic recycling significantly impaired cell polarity, migration, and also cancer stemness. Importantly, *in vivo* experiments showed that the loss of DNM1 significantly suppressed peritoneal metastatic colonization. Interestingly, metastatic cells with elevated DNM1-mediated endocytosis showed increased susceptibility to nanoparticle delivery. Collectively, these results establish the DNM1-N-cadherin axis as an important regulator of EMT-associated ovarian cancer metastasis and suggest its potential as a biomarker for targeted nanodrug therapy.

## INTRODUCTION

Epithelial-to-mesenchymal transition (EMT) is a crucial driver of metastasis, the leading cause of cancer deaths. EMT unlocks cellular plasticity by allowing cancer cells to switch from an epithelial to a more migratory, invasive, and stem-like mesenchymal state (Nieto et al., 2016; Dongre & Weinberg 2019; Yang et al., 2020). A hallmark of EMT is cadherin switching. Particularly, the upregulation of N-cadherin, rather than decreased E-cadherin, has been linked to increased metastatic signaling and behaviors (Wheelock et al., 2008; Mrozik et al., 2018). However, the precise mechanisms governing the plastic transition remain poorly understood.

Ovarian cancer is the leading cause of death among all gynecologic malignancies, mainly due to late diagnosis and widespread peritoneal metastasis. The 5-year survival rate for advanced-stage patients is less than 30% (Lheureux et al., 2019). Multiple lines of evidence have suggested the involvement of EMT in different metastatic phases of ovarian cancer within the peritoneum, a process distinct from blood-borne metastasis (Loret et al., 2019). During the initial dissemination step, EMT could promote the formation of multicellular spheroids comprised of cancer cells exfoliated from the primary site. The mesenchymal phenotype could confer increased resistance to anoikis, allowing these spheroids to survive and proliferate within the malignant ascitic fluid (Loret et al., 2019). Upon arrival at the secondary site, the mesenchymal spheroids demonstrated higher efficiency in disrupting the mesothelial cell lining covering the peritoneal organs, facilitating their implantation at distant sites (Kenny et al., 2014). To colonize the distant sites, these mesenchymal cells would undergo mesenchymal-to-epithelial transition (MET) to regain proliferative capacity (Terry et al., 2017; Dongre & Weinberg, 2019). In addition to these metastatic steps, EMT could also promote chemoresistance and immunosuppression in ovarian cancer (Loret et al., 2019; Yakubovich et al., 2023).

Targeting the EMT process represents a promising therapeutic approach to reduce cellular plasticity and metastasis in ovarian cancer, whose pathogenesis does not appear to be driven by dominant driver gene mutations, but involves more complex genomic and epigenetic alterations. Nonetheless, the clinical translation of EMT-targeting compounds has lagged behind their therapeutic potential due to the technical challenges of targeting the conventional regulatory network of EMT. This network is comprised of transcription repressors which exhibit functional redundancy and have essential roles in tissue homeostasis (Nieto et al., 2016). The search for novel, non-transcriptional regulators of EMT-associated cellular plasticity will therefore provide important therapeutic insights against metastatic diseases.

In this study, we have identified the role of DNM1, a core component of the endocytic complex, in driving ovarian cancer cells toward a plastic mesenchymal state and promoting metastatic colonization. Our results highlight the significance of endocytic recycling and N-cadherin glycosylation underlying these processes, and also suggest potential opportunities for exploiting DNM1-associated plasticity through targeted nanomedicine approaches.

## RESULTS

### DNM1 as a novel master regulator of EMT

We have developed a robust framework for identifying master regulators (MRs) in cancer development, which integrates regulons with computational tools designed to prioritize MRs and determine their functional significance. This method has been proven highly effective and powerful in identifying previously unknown oncogenes and tumor suppressors, as demonstrated by our previous data analysis with experimental validations (Ru*, et al.*, 2018; Ru *et al*., 2019). To identify potential EMT regulators, we used the hallmark epithelial and mesenchymal signatures to distinguish the subtypes of each patient in the TCGA datasets, and then compared differentially regulated genes between epithelial and mesenchymal groups. Using the regulons inferred by Algorithm for the Reconstruction of Accurate Cellular Networks (ARACNE), we identified approximately 6,700 activated EMT and 9,000 repressed EMT MRs. The workflow of our algorithm was shown in Fig. 1A. Transcription factors such as SNAI1, SNAI2, ZEB1, ZEB2, and TWIST1 were among the predicted regulators inducing EMT, validating the robustness of our approach. Further screening for non-transcription factors led to the identification of DNM1, a member of the dynamin subfamily of GTP binding proteins, as a novel master regulator (Fig. 1B and 1C). The expression of DNM1 was observed to have a negative correlation with the expression of the epithelial marker E-cadherin, but a positive correlation with the expression of the mesenchymal marker N-cadherin in TCGA ovarian cancer patient samples (Fig. 1D). Furthermore, an analysis of the four molecular subtypes of ovarian cancer (Tothill et al., 2008; The Cancer Genome Atlas Research Network, 2011) revealed that the mesenchymal subtype was associated with significantly higher expression levels of DNM1 compared to the other subtypes (Fig. 1E). A trend of higher DNM1 mRNA expression was also observed in advanced-stage (III/IV) ovarian cancer patients compared to those in early stages (I/II) by qPCR (Fig. 1F). DNM1 mRNA expression was higher in ovarian tumor tissue compared to normal tissue in the TCGA dataset, and high DNM1 expression was associated with progression-free survival and post-progression survival (Supplementary Fig. 1A and 1B). In contrast, DNM2 expression did not exhibit a significant difference between tumor and normal tissue, while DNM3 showed lower expression in tumor tissue compared to normal tissue (Supplementary Fig. 1A). We further confirmed DNM1 expression in a panel of ovarian cancer cell lines characterized by different EMT states (Huang et al., 2013; Tan et al, 2014; Terraneo et al., 2020). Higher DNM1 expression was shown in intermediate epithelial (OVCA429), intermediate mesenchymal (Kuramochi), and mesenchymal ovarian cell lines (HEYA8, A2780) compared to epithelial lines (OVSAHO, CaOV3 and OVCAR3) (Fig. 1G). These data together support a strong association between DNM1 and EMT in ovarian cancer (Fig. 1G).

**Figure 1.**
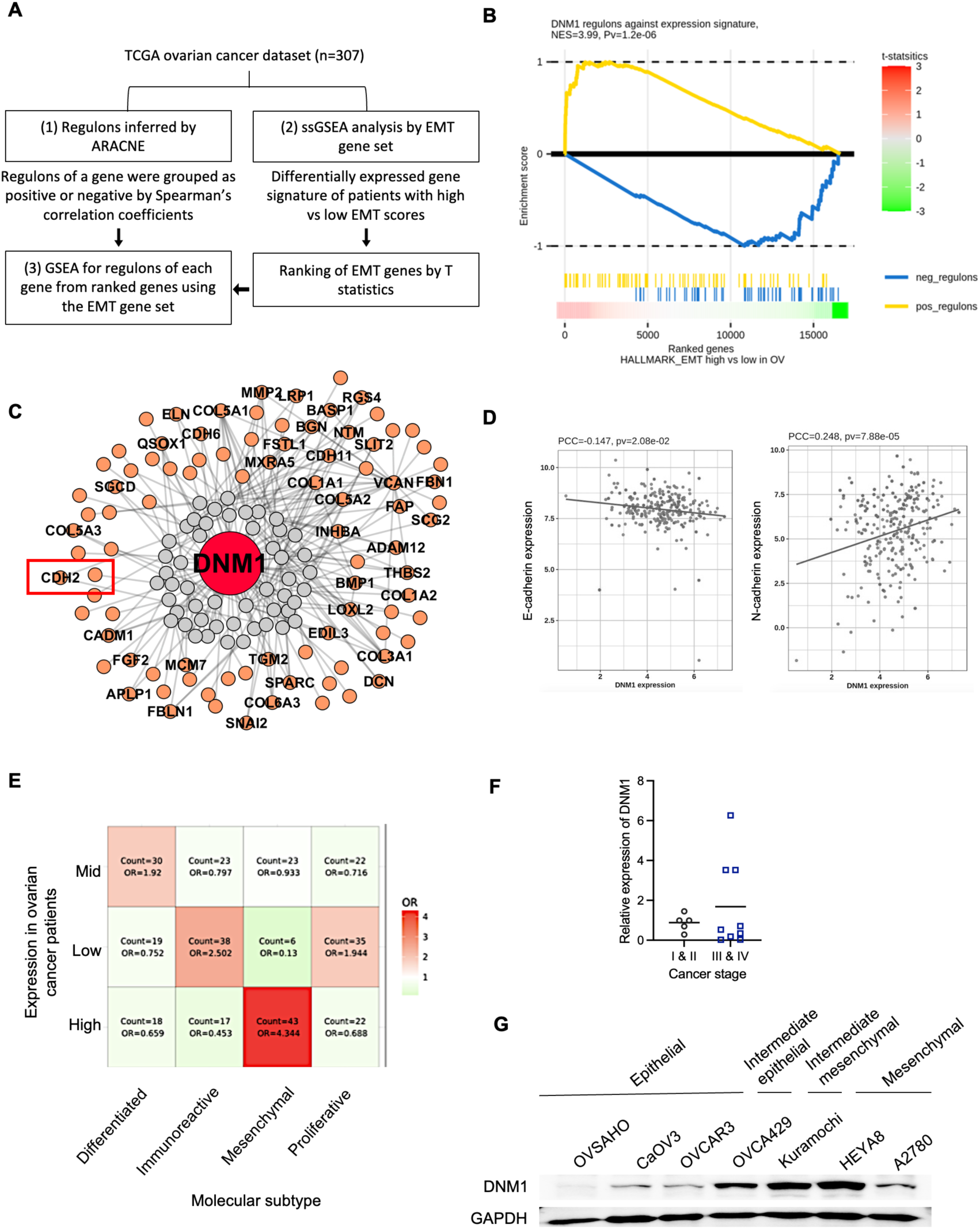
DNM1 is identified as a EMT master regulator in ovarian cancer patients. (**A**) A master regulator (MR) algorithm integrating The Cancer Genome Atlas (TCGA) ovarian cancer dataset and EMT signature was used to infer potential EMT markers. The basic workflow was shown: (1) Based on the ovarian cancer gene expression profile from TCGA dataset, transcriptional targets (or regulons) were inferred by using Algorithm for the Reconstruction of Accurate Cellular Networks (ARACNE) with default parameters. The regulons of a potential MR were divided into positive (upregulated) and negative (downregulated) groups based on the Spearman’s correlation coefficients between the MR expression level and each gene in its regulon; (2) Differentially expressed gene (DEG) test was performed to compare EMT high tumors vs EMT low tumors. EMT scores were calculated using single sample gene set enrichment analysis (ssGSEA) methods from the HALLMARK M gene set to get a ranked differential gene signature (by T statistics); (3) The ranked gene list was then used as input for the GSEA method from the R “gage” package to evaluate the enrichment of the regulons and their associated MRs based on FDR-adjusted P values. Only MRs with significantly enriched associated regulons (cut-off < 0.05) were considered for further experiment validations. (**B**) GSEA analysis revealed a significant correlation between DNM1 and HALLMARK_EPITHELIAL_MESENCHYMAL_TRANSITION. (**C**) A gene regulatory network between DNM1 and its associated regulons from HALLMARK_EPITHELIAL_ MESENCHYMAL_TRANSITION were generated using ARACNE. (**D**) Negative correlation between DNM1 and E-cadherin transcript levels (left panel) and positive correlation between DNM1 and N-cadherin levels (right panel) were shown. (**E**) Odd ratio (OR) of high, mid-, or low DNM1 expression in different molecular subtypes of ovarian cancer were calculated. (**F**) Real-time PCR was performed for DNM1 in early (I/II) and late (III/IV) stage high grade serous ovarian cancer patient samples. (**G**) DNM1 expression in ovarian cancer cells with different EMT phenotypes were analyzed by Western blot. GAPDH served as a loading control.

### DNM1 promotes metastatic competence

To investigate the functional significance of DNM1 in EMT, we utilized an isogenic model, in which the highly metastatic (HM) cells, unlike their non-metastatic (NM) counterparts, exhibited a specific capability for migration and peritoneal metastasis (To et al., 2017). We found that HM cells had higher expression of DNM1 and displayed a mesenchymal-like phenotype with higher expression of N-cadherin and vimentin, along with little or no expression of the epithelial marker E-cadherin, compared to NM cells (Fig. 2A). Silencing DNM1 led to significant inhibition of the mesenchymal morphology, as well as a decrease in the migration ability of HM cells by ∼1.47-fold (Fig. 2B and 2C). Interestingly, DNM1-mediated migration was independent of the well-known EMT inducer TGF-β (Supplementary Fig. 2A). Specific knockdown of DNM1 by siRNA was confirmed by Western blot (Supplementary Fig. 2B). Conversely, overexpression of DNM1 in NM cells promoted EMT phenotypes, resulting in a spindle-like morphology and increased migration by ∼1.5-fold (Fig. 2B and 2D). The MTT assay showed that DNM1 had little effect on the growth rate of ovarian cancer cells, suggesting that its migratory/invasive effects are independent of cell growth (Supplementary Fig. 2C). We also treated the cells with dynasore, a GTPase inhibitor of dynamin-mediated endocytosis (Kirchhausen et al., 2008), and found that treatment of dynasore significantly reduced cell migration (Fig. 2E). Cancer stem-like phenotypes are a manifestation of EMT plasticity, and DNM1 silencing reduced both the size and number of spheroids in sphere formation assays (Fig. 2F). To further confirm the *in vivo* role of DNM1, luciferase-expressing HM cells stably transfected with non-specific (NS) or DNM1-specific shRNA were intraperitoneally injected into female NOD/SCID mice. Bioluminescence imaging revealed that mice injected with DNM1 shRNA cells exhibited significantly less peritoneal dissemination and higher metastatic tumor burdens in the mesenteries (∼2.04-fold) compared to mice injected with NS shRNA (Fig. 2G and 2H). These findings suggest that DNM1 not only promotes initial spreading but also promotes metastatic colonization.

**Figure 2.**
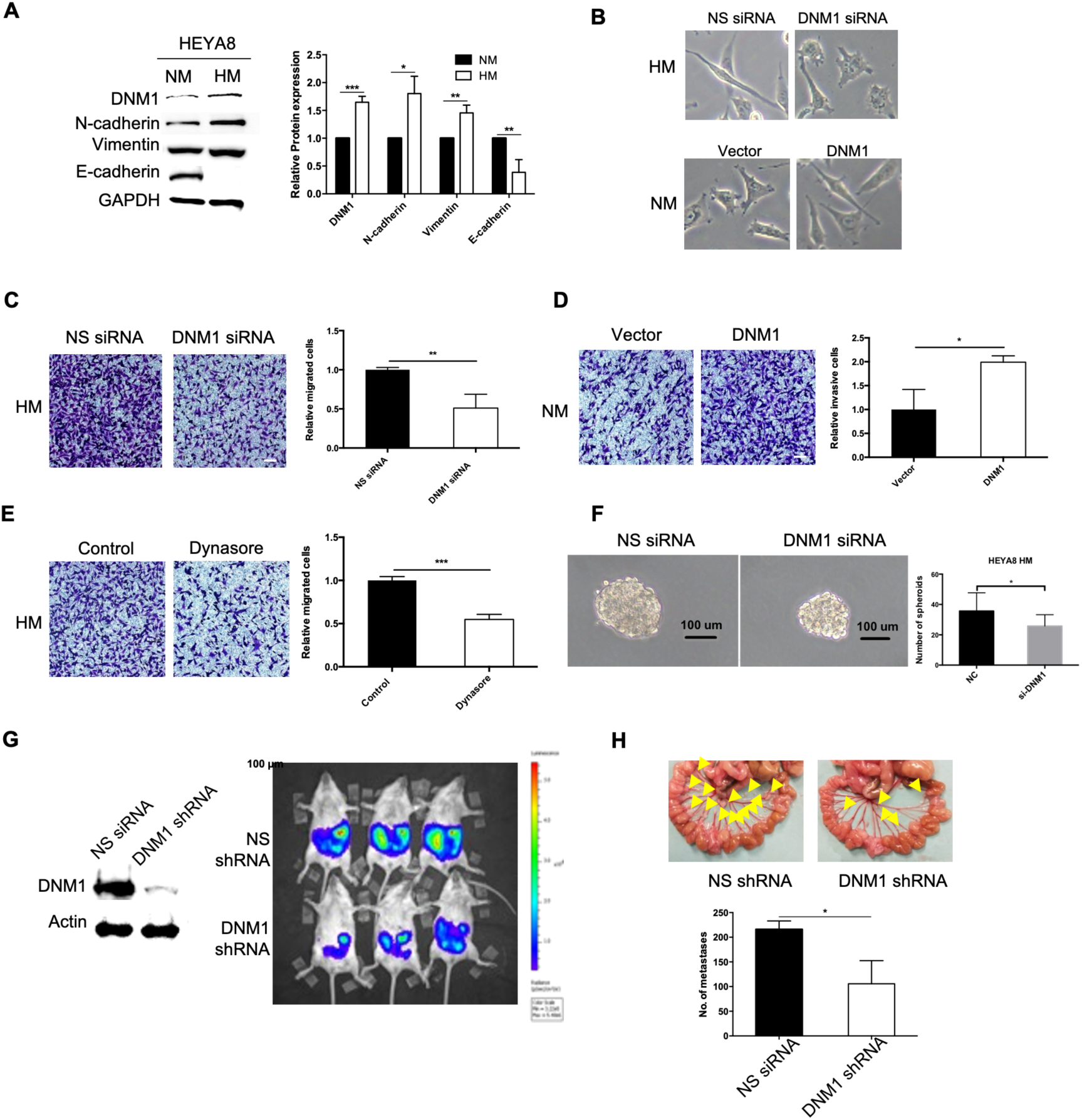
DNM1 enhances migration and metastasis. (**A**) Expression levels of DNM1, N-cadherin, vimentin, E-cadherin were analyzed in NM and HM cells. (**B**) Morphologies of HM cells transfected with non-specific (NS) or DNM1 siRNA, and NM cells transfected with control vector or DNM1 overexpression vector, were observed under a light microscope. (**C**) HM cells were transfected with NS or DNM1 siRNA, followed by migration assays. Migrated cells were stained with crystal violet and counted in five random fields of view. (**D**) NM cells were transfected with control vector or DNM1 overexpression vector, followed by migration assays. Migrated cells were stained with crystal violet and counted in five random fields of view. (**E**) HM cells were treated with vehicle control or dynasore (100 µM) and subjected to migration assays. Migrated cells were stained with crystal violet and counted. (**F**) HM cells were transfected with NS or DNM1 siRNA, followed by sphere formation assays. Representative spheroids were shown and the number of spheres were counted. (**G**) DNM1 expression was tested in HM cells that were stably transfected with NS or DNM1 shRNA. These cells were intraperitoneally injected into NOD-SCID mice (n=3 per group, repeated twice). Peritoneal metastasis was visualized by in vivo bioluminescence imaging. (**H**) Metastatic nodules at the mesenteries were counted after the mice were scarified. Band intensities were quantified by ImageJ, and GAPDH served as a loading control for all Western blot analyses. Data shown in A, C-F are representative of three independent experiments. *, *P* < 0.05. **, P < 0.01 and ***, *P*< 0.005.

### DNM1 enhances N-cadherin levels via endocytosis and subsequent recycling to the membrane

To investigate the effect of DNM1 manipulation on EMT markers, we probed for E-cadherin, N-cadherin, and vimentin in cells with DNM1 knockdown or overexpression. Silencing of DNM1 consistently inhibited the expression of N-cadherin and vimentin in HM, Kuramochi, and OVCA429 cells (Fig. 3A, 3C and 3D), whereas overexpression of DNM1 enhanced N-cadherin and vimentin levels in NM and CaOV3 cells (Fig. 3B and 3E) with little or no changes in E-cadherin in these cells (Fig. 3A-E). Based on these observations and given that N-cadherin is one of the regulons of DNM1 (Fig. 1C), we speculate that DNM1 could regulate EMT via N-cadherin. Consistently, we observed a reduction of N-cadherin in mice tumors with DNM1 shRNA compared to NS shRNA control (Fig. 3F).

**Figure 3.**
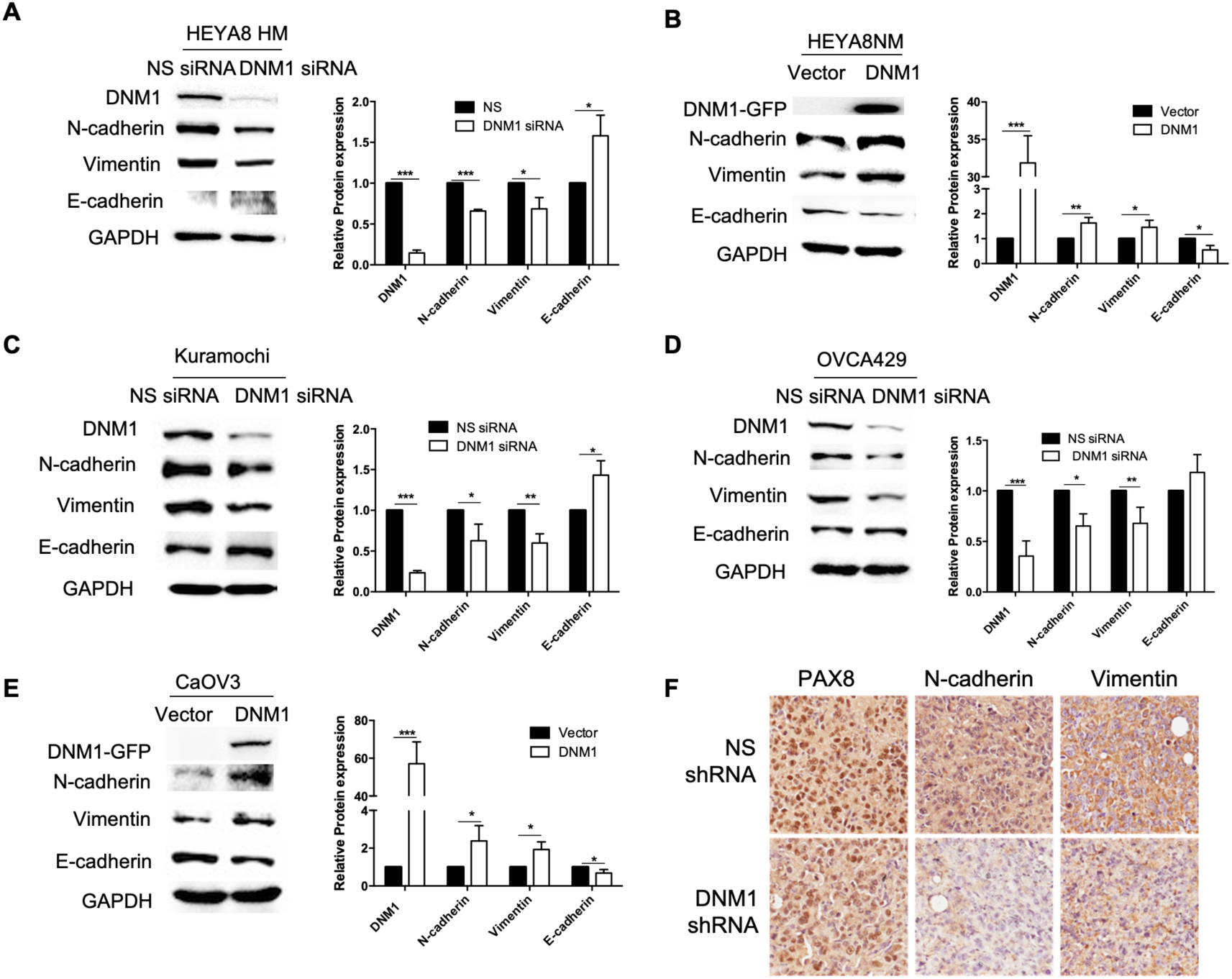
DNM1 regulates N-cadherin expression. Protein expression of DNM1, N-cadherin, vimentin, and E-cadherin were analyzed in **(A)** HM cells transfected with NS or DNM1 siRNA, **(B)** NM cells transfected with control vector or DNM1 overexpression vector, **(C)** Kuramochi cells transfected with NS or DNM1 siRNA; **(D)** OVCA429 cells transfected with NS or DNM1 siRNA; and (E) CaOV3 cells transfected with control vector or DNM1 overexpression vector. Band intensities were quantified by imageJ, and GAPDH served as a loading control for all Western blot analyses. Representative blots of three independent experiments were shown. *, *P* < 0.05. **, *P* < 0.01 and ***, *P* < 0.005. **(F)** Immunohistochemistry of PAX8 (tumor marker), N-cadherin and vimentin were performed in metastatic tumors in mice formed by HM with NS or DNM1 shRNA.

DNM1 belongs to the dynamin subfamily of GTP-binding proteins, which possesses unique mechanochemical properties involved in vesicle scission during endocytosis and vesicular trafficking (Meng et al., 2017). To investigate whether DNM1 could alter N-cadherin turnover, we conduct ed cell surface biotinylation and internalization assays as illustrated in Fig. 4A. Silencing of DNM1 significantly inhibited the endocytosis of N-cadherin (Fig. 4B). Cadherin protein turnover is regulated by two major cellular uptake mechanisms, clathrin- and caveolae-mediated endocytosis (Kowalczyk & Nanes, 2012). Therefore, we examined whether these mechanisms are involved in DNM1-mediated N-cadherin endocytosis. Silencing of caveloin-1 (CAV1) inhibited N-cadherin endocytosis (Fig. 4F), whereas silencing of clathrin (CLTC) had no effect (Fig. 4G). Knockdown of CAV1 and CLTC was confirmed by Western blot (Supplementary Fig. 2D-E). These results indicate that DNM1 may mediate N-cadherin endocytosis via a caveolae-dependent rather than a clathrin-dependent mechanism.

**Figure 4.**
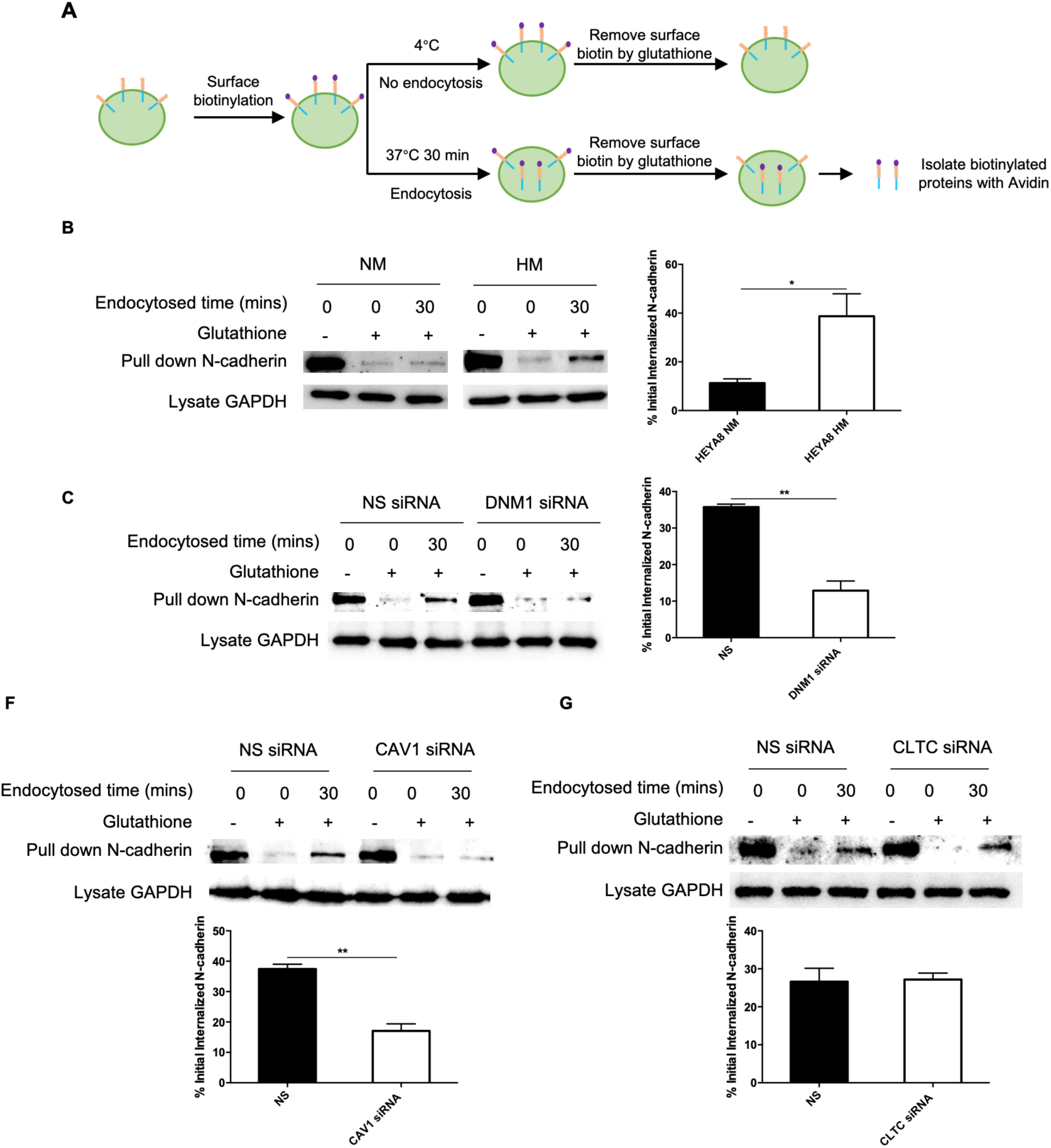
DNM1 enhances N-cadherin levels through caveolae-mediated endocytosis. **(A)** The simplified workflow of the endocytosis assay using biotin-labeling was shown. Briefly, surface proteins were biotinylated, followed by incubation at 4°C as control or at 37°C to allow for endocytosis. Surface biotin was then removed by incubating with a glutathione solution. After quenching of free glutathione, cells were lysed and biotinylated proteins were isolated by avidin. Endocytosis of N-cadherin was compared between **(B)** NM and HM cells, **(C)** HM cells transfected with NS or DNM1 siRNA, **(D)** HM cells transfected with NS or caveolin-1 (CAV1) siRNA, and **(E)** HM cells transfected with NS or clathrin (CLTC) siRNA using the biotin-labelling assay, followed by Western blot. Band intensities were quantified by ImageJ, and GAPDH of total cell lysate served as a loading control. Representative data of three independent experiments were shown. *, *P* < 0.05. **, *P* < 0.01 and ***, *P* < 0.005.

Once membrane proteins are endocytosed, they can either be sorted to the lysosome for degradation or to recycling endosomes for recycling back to the cell surface. Therefore, we investigated the role of DNM1 in N-cadherin recycling using biotin-labeling assays (Fig. 5A). Silencing of DNM1 in HM cells inhibited N-cadherin recycling, while overexpression of DNM1 in NM cells enhanced N-cadherin recycling (Fig. 5B and 5C). Consistently, treatment with a proteasome inhibitor, MG132, inhibited N-cadherin reduction caused by DNM1 knockdown, suggesting that DNM1 could divert N-cadherin from degradation to the endosomal recycling pathway (Supplementary Fig. 2D). In addition, we observed decreased colocalization between DNM1 and Rab11, a recycling endosome marker, in HM cells treated with DNM1 siRNA, whereas increased colocalization was observed in NM cells overexpressing DNM1 (Fig. 6D and 6E), confirming a role for DNM1 in N-cadherin endocytic recycling in ovarian cancer cells. As recycling endosomes play a central role in the regulation of cell polarity (Eaton et al., 2014), we assessed the role of DNM1 in cell migration using scrape wounding assays followed by Golgi tracking. We found that DNM1 siRNA treated HM cells exhibited significantly impaired Golgi positioning towards the wound, as well as reduced directional recycling of N-cadherin-containing vesicles to the cell membrane at the leading edge (Fig. 5F and 5G). Similar results were observed in Kuramochi cells (Supplementary Fig. 3A and 3B). We also used primaquine, an inhibitor of the endosomal recycling pathway (van Weert et al., 2000), and found that it reduced total level of N-cadherin and its recycling (Fig. 6A-B, Supplementary Fig. 3C). Primaquine treatment also reduced cell migration, impaired both Golgi polarization, and reduced the recycling of N-cadherin back to the cell membrane at the leading edge (Fig. 7C-D; Supplementary Fig. 3D-F). Taken together, these results indicate that N-cadherin is endocytosed and recycled in a DNM1-dependent manner for directional cell migration.

**Figure 5.**
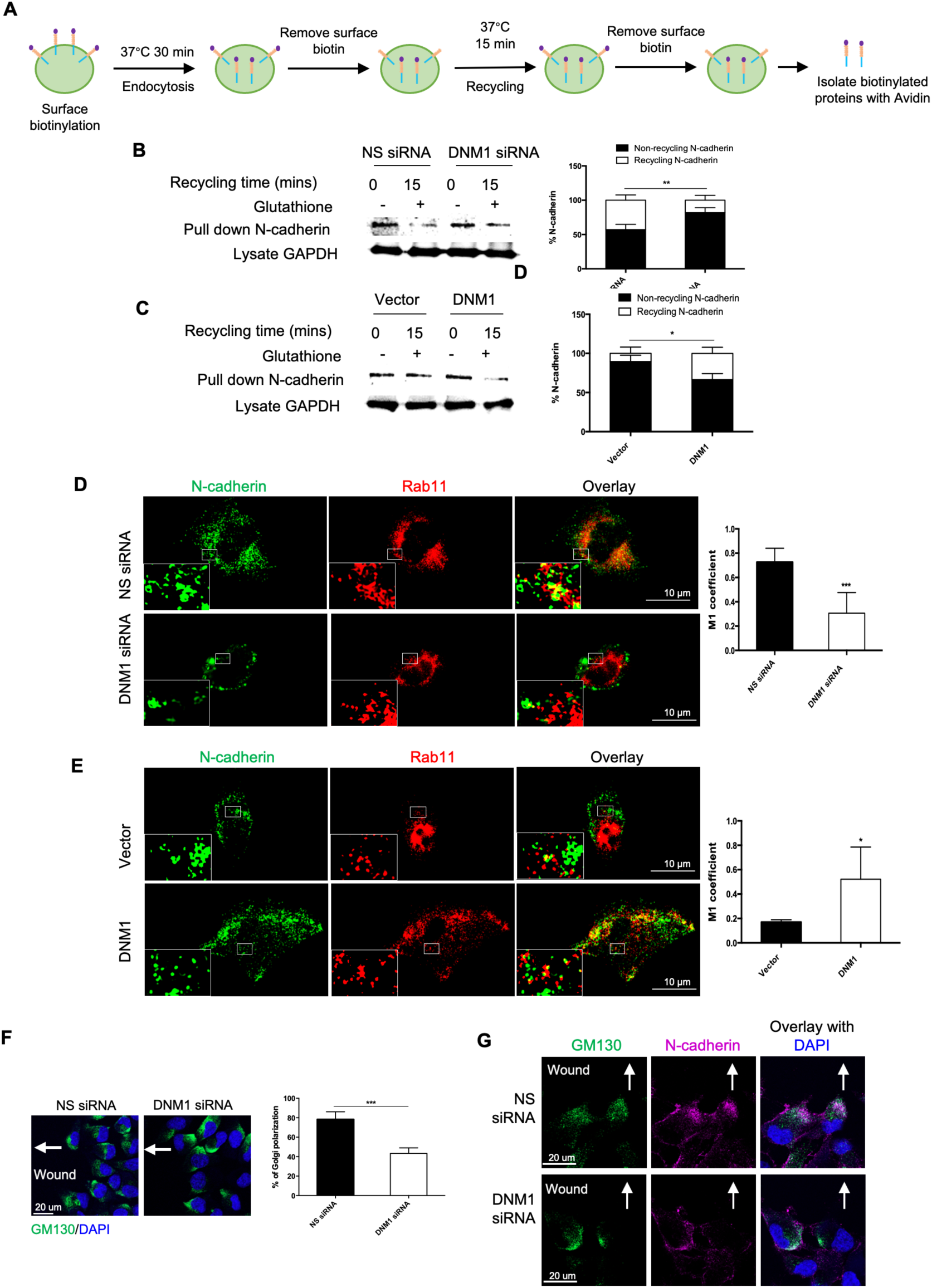
DNM1 enhances recycling of N-cadherin. **(A)** The simplified workflow of recycling assay using biotin-labeling was shown. Briefly, surface proteins were biotinylated, followed by incubation at 37°C to allow for endocytosis. Surface biotin was then removed by glutathione. Cells were further incubated at 37°C to allow for recycling. Surface biotin was again removed by glutathione. Cells were lysed and the remaining biotinylated proteins, which were not recycled, were isolated by avidin. Recycling of N-cadherin was compared between **(B)** HM cells transfected with NS or DNM1 siRNA, and **(C)** NM cells transfected with control or DNM1 overexpression vectors, using the biotin-labelling assay, followed by Western blot. Band intensities were quantified by ImageJ. GAPDH of total cell lysate served as a loading control. **(D)** HM cells transfected with NS or DNM1 siRNA were stained for N-cadherin (green) and Rab11 (marker of recycling endosomes, red). Colocalization of the two proteins was observed by confocal imaging. **(E)** NM transfected with control or DNM1 overexpression vector were stained for N-cadherin (green) and Rab11 (red). Colocalization of the two proteins was observed by confocal imaging. **(F)** Cell-free gaps were created in HM monolayers after transfection with NS or DNM1 siRNA. Golgi orientation was visualized by staining the Golgi marker GM130 (green). The number of cells with correct polarity were counted. **(G)** The indicated cells were co-stained with GM130 and N-cadherin (magenta), followed by observation by confocal imaging. Cell nuclei were counterstained by DAPI. Arrows indicated the expected Golgi direction towards the wound. Representative data of 3 independent experiments were shown. *, *P* < 0.05. **, *P* < 0.01 and ***, *P* < 0.005.

**Figure 6.**
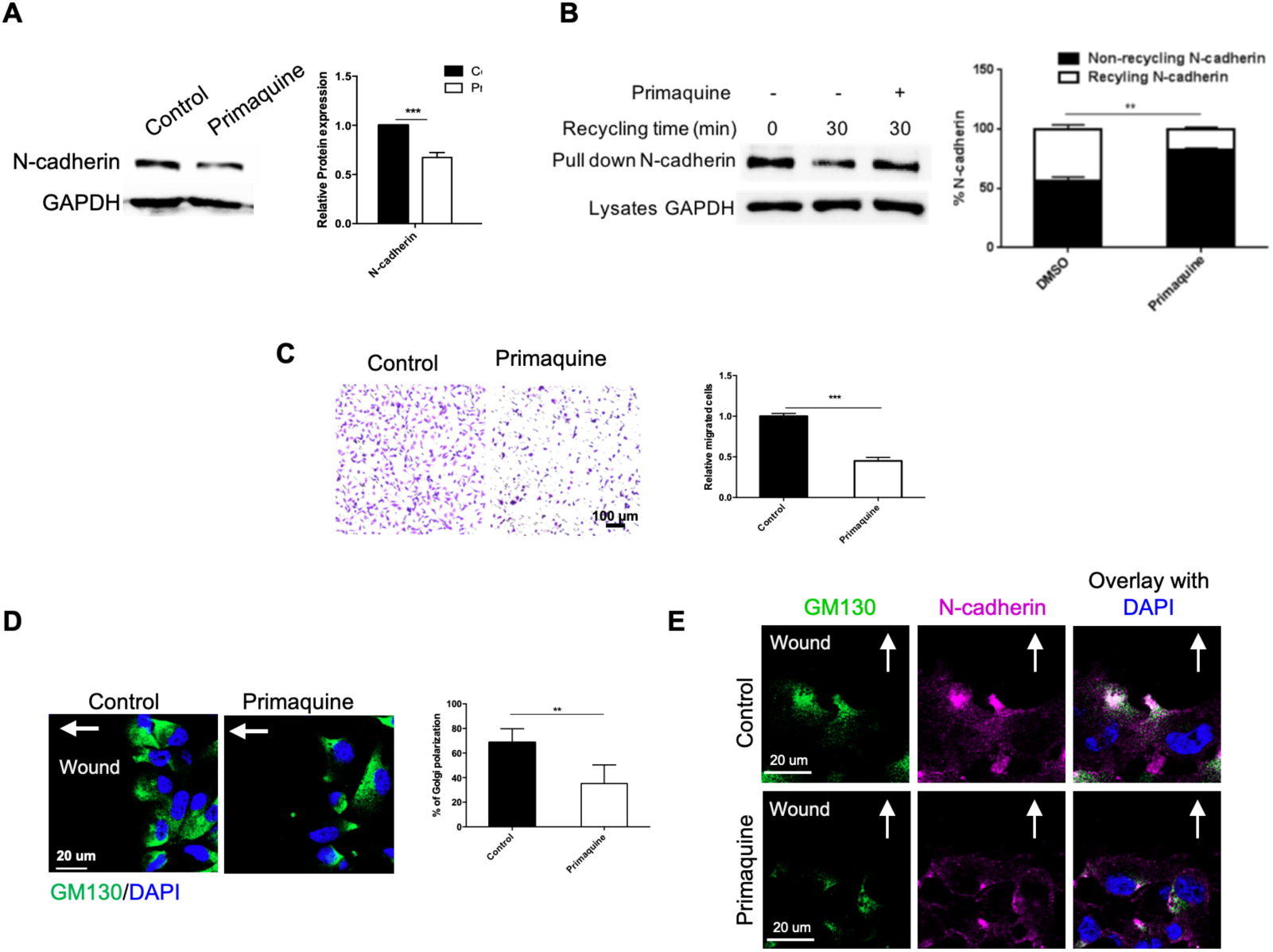
Treatment with recycling inhibitor reduces N-cadherin level, migration, and polarization. HM cells were treated with vehicle control or primaquine (100 μM), a recycling inhibitor, and analyzed by the following assays. **(A)** N-cadherin expression was tested by Western blot. **(B)** Biotin recycling assay was performed, followed by Western blot for N-cadherin. **(C)** Migration assays were performed. Migrated cells were stained with crystal violet and counted in five random fields of view. **(D)** Cell-free gaps were created in the cell monolayer. Golgi orientation was visualized by staining the Golgi marker GM130 (green). The number of cells with correct polarity were counted. **(E)** The indicated cells were co-stained with GM130 and N-cadherin (magenta), followed by confocal imaging. Nuclei were counterstained by DAPI (blue). Arrows indicated the expected Golgi direction towards the wound. Representative data of three independent experiments were shown. *, *P* < 0.05. **, *P* < 0.01 and ***, *P* < 0.005.

**Figure 7.**
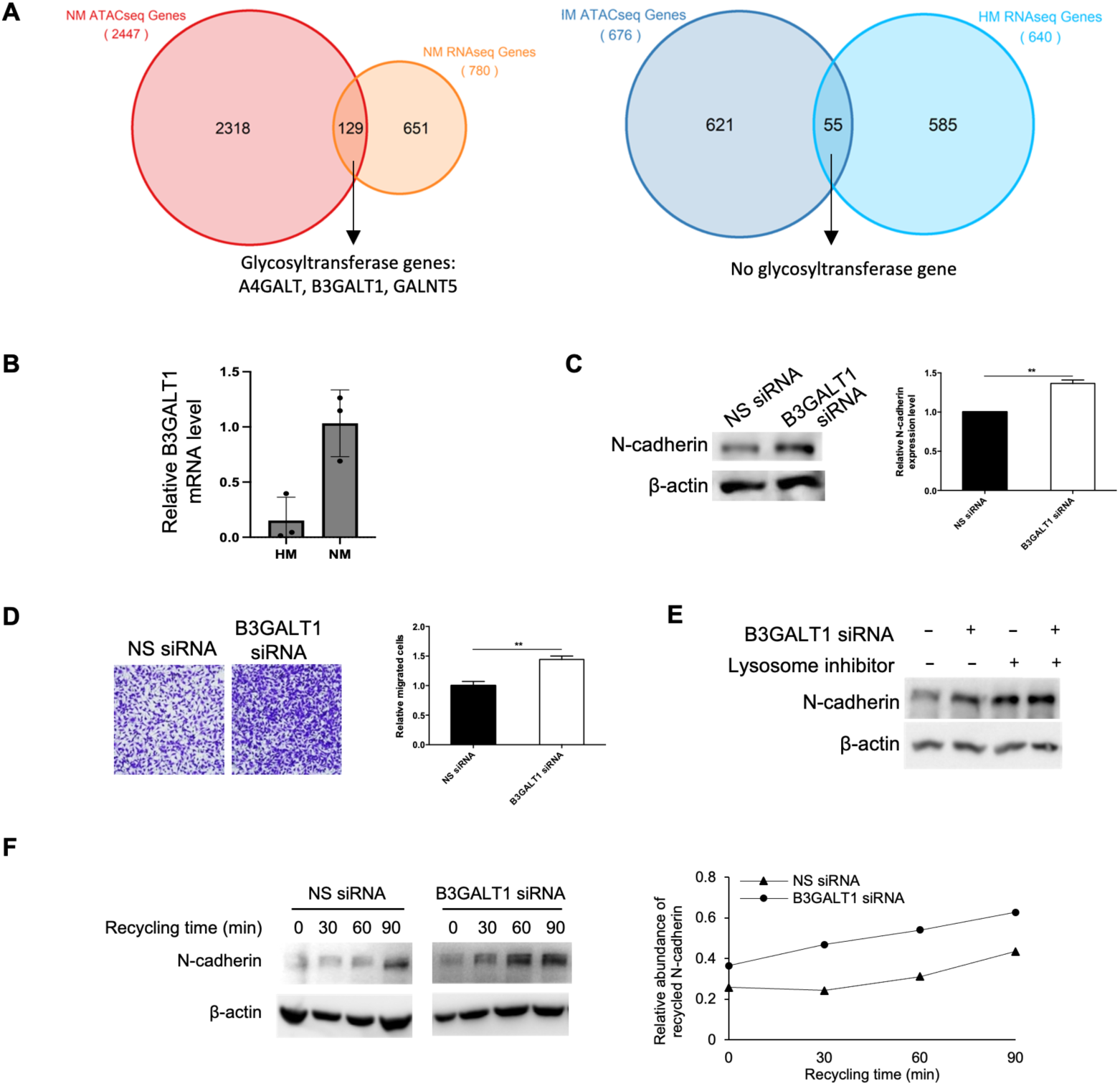
B3GALT1-mediated glycosylation inhibits N-cadherin recycling by DNM1. **(A)** ATAC-seq and RNA-seq were performed with NM and HM cells (3 replicates each). Differentially accessible peak-associated genes were overlapped with upregulated genes in NM or HM cells. **(B)** Higher expression of B3GALT1 in NM compared to HM was confirmed by qPCR. Actin was used as a loading control. **(C)** N-cadherin expression was tested by Western blot in NM cells transfected with NS or B3GALT1 siRNA. **(D)** Migration assays were performed after transfection of NS or B3GALT1 siRNA in NM cells. Migrated cells were stained with crystal violet and counted in five random fields of view. **(E)** NM cells were transfected with NS or B3GALT1 siRNA, and treated with or without 50 mM NH_4_Cl (lysosome inhibitor) for 4 h. N-cadherin expression was detected by Western blot. **(F)** NM cells were transfected with NS or B3GALT1 siRNA. After calcium chelation to induce N-cadherin internalization, cell surface proteins were allowed to recover for the indicated time points. Cell surface proteins were biotinylated and the recycled N-cadherin was analysed by Western Blot. Band intensities were quantified by ImageJ, and β-actin of total cell lysate served as a loading control. Representative data of three independent experiments were shown. *, *P* < 0.05. **, *P* < 0.01 and ***, *P* < 0.005.

### B3GALT1 regulates N-cadherin endocytic recycling

N-glycosylation is a common protein modification that plays a crucial role in regulating rapid, dynamic cadherin function (Carvalho et al., 2016). Using ATAC-seq combined with RNA-seq analysis (Supplementary Fig. 5A), we identified three glycosyltransferase genes, namely A4GALT, B3GALT1, and GALNT5, which exhibited increased chromatin accessibility and gene expression in NM cells compared to HM cells (Fig. 7A). Among these genes, A4GALT1 has been shown to inhibit EMT in ovarian cancer by modulating glycosphingolipids (Jacob et al., 2018), while the roles of different GALNT members in EMT have also been implicated (Beaman et al., 2022). Therefore, we focused on B3GALT1, which has an unknown role in EMT. B3GALT1 was upregulated in NM cells compared to HM cells, as confirmed by qPCR (Fig. 7B). Knockdown of B3GALT1 increased N-cadherin levels and enhanced cell migration ability in NM cells (Fig. 7C). Similar increase in N-cadherin and cell migration were observed in Kuramochi cells transfected with B3GALT1 siRNA (Supplementary Fig. 4E). Treatment with ammonium chloride (NH_4_Cl), a lysosome inhibitor. restored N-cadherin levels, indicating that N-cadherin undergoes continuous lysosomal degradation in NM cells (Fig. 7E). Interestingly, B3GALT1 knockdown further increased N-cadherin protein levels in lysosome-inhibited NM cells, suggesting that B3GALT1 can reduce N-cadherin stability independent of lysosomal degradation (Fig. 7E). Moreover, B3GALT1 knockdown accelerated the recycling of N-cadherin in NM cells (Fig. 7F). These findings suggest that B3GALT1 may function as an EMT suppressor by reducing N-cadherin glycosylation and recycling.

### DNM1 enhances the susceptibility of metastatic cells to nanoparticle uptake for targeted drug delivery

We have previously developed self-assembling supramolecular dendrimers as precision nanomaterials for the delivery of imaging agents, small molecular anticancer drugs, and nucleic acid therapeutics for cancer detection and treatment (Lyu et al., 2020; Chen et al., 2023; Jiang et al., 2023) (Fig. 8A). Nanoparticles exploit various endocytic pathways for cellular entry, and caveolae-dependent endocytosis is the preferred one for drug delivery as it prevents massive integration into lysosomes for enzymatic degradation while promoting effective cargo release from endosomes (Hillaireau & Couvreur, 2009; Blanco et al., 2015; Du Rietz et al., 2020). In this study, we investigated the potential therapeutic effectiveness of harnessing DNM1-mediated endocytosis for our dendrimer nanotechnology-based drug delivery. Using dendriplexes containing fluorescein-labeled siRNA, we found that HM cells are more capable of nanoparticles uptake as compared to NM cells (Fig. 8B). DNM1 knockdown significantly reduces the uptake by HM cells (Fig. 8C). These data suggest that DNM1 enables metastatic cells for preferential uptake of nanoparticles, making targeted drug delivery more favorable and effective.

**Figure 8.**
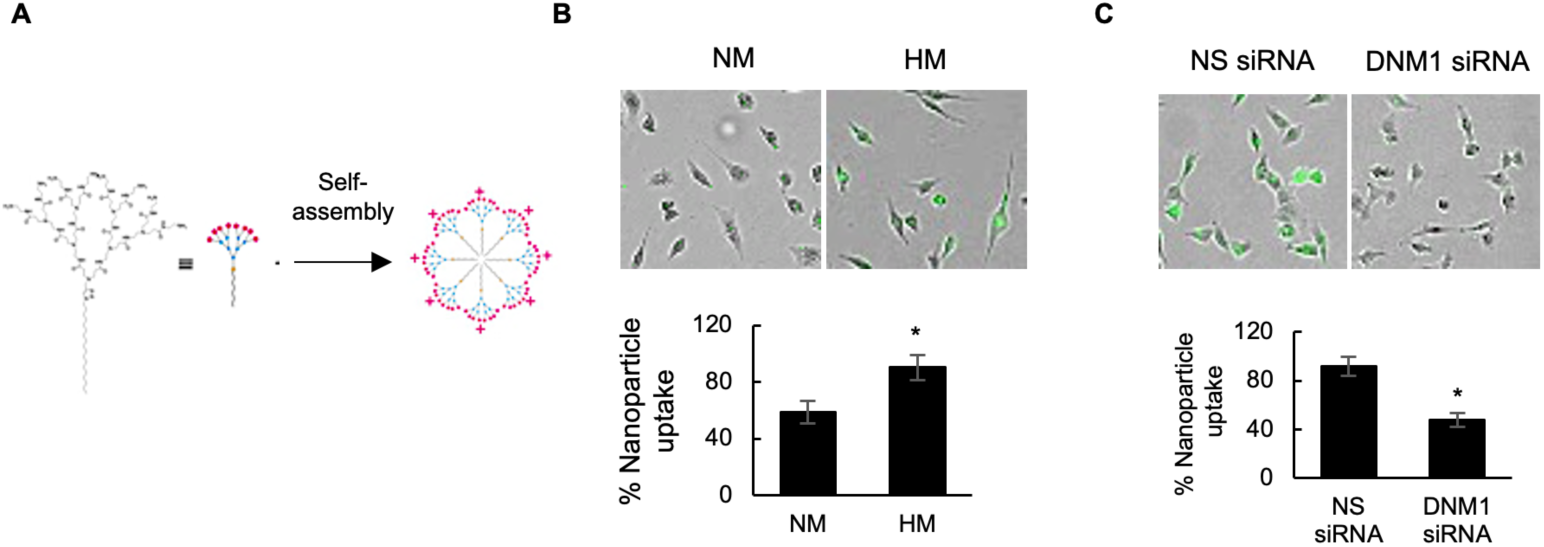
DNM1 enhances endocytosis of nanoparticle-encapsulated siRNA by metastatic cells. **(A)** Schematic diagram of self-assembly of nanoparticle into micelles for endocytic uptake. **(B)** NM and HM cells were treated with nanoparticles containing fluorescein-labeled siRNA (green) at 20 nM. **(C)** HM cells were transfected with NS or DNM1 siRNA, followed by treatment with nanoparticles containing fluorescein-labeled siRNA (green) at 20 nM. The percentage of cells taken up the dendriplexes was quantified by counting the number of cells giving fluorescent singles normalized by the total cell number in the field. Data are represented as mean ± SD, *, *P* < 0.05.

## DISCUSSION

EMT is a key driver of metastasis and therapy resistance, rendering it a promising therapeutic target. However, the direct pharmacological inhibition of EMT-inducing transcription factors has been challenging. Hence, identifying novel transcription-independent regulators is critical for developing more effective treatment strategies. In this study, using our established bioinformatic algorithms to analyze multiple TCGA datasets, we identified DNM1 as a novel master regulator of EMT which drives ovarian cancer metastasis independent of transcriptional regulation.

Exogenous EMT-inducing signals, such as TGF-β, often employ receptor-mediated endocytic pathways to promote mesenchymal transformation (Corallino et al., 2015). However, how intrinsic signaling pathways, particularly the endocytic circuitry, could autonomously control EMT plasticity remains unclear. Moreover, while the endocytic degradation of cadherin molecules is extensively studied (Corallino et al., 2015), the mechanistic basis for their endocytic recycling remains poorly understood. Here, we found a previously unrecognized role of DNM1 in regulating N-cadherin recycling independent of TGF-β to maintain the plastic mesenchymal feature during metastasis. Our data indicate that DNM1 regulates N-cadherin in a caveolae-dependent manner, instead of through its reported role in clathrin-mediated endocytosis (Meng, 2017). Relevant to our findings, the dynamic endosomal trafficking of cadherins has been suggested to offer more physiologically relevant control over EMT-associated cancer cell motility than the conventional transcriptional repression program (Aiello et al., 2018). Notably, DNM1 was shown to mediate the ultrafast endocytosis of synaptic vesicles in neurons (Imoto et al., 2017), which is much faster than the conventional clathrin-mediated endocytosis. Such rapid internalization would probably facilitate and sustain the spatial reorganization of key signaling cascades within cancer cells. While some reports have suggested that EMT may not be strictly necessary for metastasis in certain mouse models, these studies have mainly relied on lineage tracing using gene reporters or genetic ablation of major EMT transcription factors like Snail or Twist (Fischer et al., 2015; Zheng et al., 2015). Such approaches have not fully captured the role of rapid endocytic mechanisms, and our identification of the DNM1-N-cadherin axis highlights the need for a more comprehensive understanding of the diverse mechanisms underlying the metastatic cascade, beyond the classical EMT transcription factors.

The delicate balance between epithelial E-cadherin and mesenchymal N-cadherin expression is crucial for EMT plasticity. Whereas E-cadherin has been the primary focus in cancer research, the significance of N-cadherin has often been overlooked, despite evidence that N-cadherin could exert a dominant influence over E-cadherin by contributing to both adhesive properties and pro-metastatic behaviors in cancer cells (Wheelok et al., 2008; Mrozik et al., 2018; Li et al., 2020). N-cadherin was shown to facilitate the formation of multicellular aggregates and their invasion in lung and ovarian cancer models (Klymenko et al. 2017; Kuriyama et al., 2016), and its polarized recycling from the cell rear to the leading edge would enhance collective migration, a characteristic feature of highly plastic EMT cells (Peglion et al., 2014, Lüönd et al., 2021). Targeting N-cadherin with a monoclonal antibody has also shown promising results in reducing prostate cancer metastasis in mice (Tanaka et al., 2010). Interestingly, we found that post-translational glycosylation affects the endocytic recycling of N-cadherin, which may provide valuable insights for more precise targeting of N-cadherin.

Various clinical data have implicated the close association between EMT and ovarian cancer progression and therapy resistance. Whereas high expression of E-cadherin is associated with better patient survival and reduced invasiveness, elevated levels of N-cadherin have been correlated with worse clinical outcomes (Veatch et al., 1994; Klymenko et al., 2017; Assidi, 2022). Multiple studies have suggested that the mesenchymal molecular subtype is associated with severe post-operative complications and also poor clinical outcomes and in ovarian cancer patients (Tothill et al., 2008; Yang et al., 2013; Vargas et al., 2015, Wang et al., 2017; Torres et al., 2018). While recent data suggests that some degree of epithelial phenotypes could be important for aggressive tumor behaviors, there have been substantial evidence that mesenchymal features are a necessary prerequisite for tumor plasticity (Cook & Vanderhyden, 2023; Wang & Yan, 2024). A hysteretic model with persistence EMT has also been proposed for metastases formation (Celià-Terrassa et al., 2020). Moreover, strong correlations between mesenchymal phenotypes and increased therapy resistance, which contributes to tumor recurrence, have been observed by different studies (Loret et al., 2021; Shi et al., 2023). A recent EMT trajectory mapping of 7,180 epithelial tumors has suggested that cancer cells harboring partial or fully mesenchymal EMT phenotypes exhibited significantly worse overall survival compared to those with a fully epithelial cellular state (Malagoli et al., 2023). Intriguingly, we have detected increased expression of the DNM1 in ovarian cancer cell lines displaying more pronounced mesenchymal phenotypes, which is supportive of the role of DNM1 in self-reinforcing the mesenchymal state of ovarian cancer cells, leading to the functional consequence of enhanced metastasis.

The findings from this study hold two significant therapeutic implications. First, the dysregulated endocytic recycling pathway represent a potential therapeutic target selective for plastic cells, as we have shown that its pharmacological inhibition impaired cellular polarization and migration. Indeed, temporary and reversible targeting of the endocytic machinery has shown promise as a strategy to increase efficacy of other anti-cancer agents such as monoclonal antibody, without causing significant side effects (Banushi et al., 2023). Second, we discovered that the enhanced endocytic capacity driven by high DNM1 expression in ovarian cancer cells may facilitate the efficient internalization of therapeutic nanoparticles. This suggests that although EMT plasticity is known to prime cancer with inherent resistance to conventional chemotherapy and targeted therapies in the highly aggressive tumor cells, it could on the other hand render them more sensitive to nanoparticle-based interventions. One key advantage of dendrimer nanocarriers is their ability to leverage the enhanced permeation and retention effect, which allows them to preferentially accumulate and deliver therapeutic agents directly to tumor lesions, thereby improving the specificity of drug targeting. However, the lack of reliable patient stratification biomarkers has been one key challenge for the successful clinical translation of cancer nanomedicine (Huang et al., 2022). Patient selection based on DNM1 expression may help identify those most likely to benefit from nanodrug formulations.

Taken together, we have uncovered a novel role for the DNM1 in regulating the endocytic recycling of the mesenchymal N-cadherin, which may generate a pool of highly plastic EMT cells capable of driving ovarian cancer metastasis. Targeting the tumor-specific EMT-associated endocytic recycling machinery may therefore be a more feasible approach than inhibiting the EMT transcriptional program. Exploring the unique vulnerabilities of the DNM1-mediated endocytic capacity could lead to the development of more effective treatment strategies against highly plastic, treatment-resistant metastatic cells.

## MATERIALS AND METHODS

### Master regulator prediction

Based on the ovarian cancer gene expression profile from The Cancer Genome Atlas Program (TCGA) dataset, transcriptional targets (or regulons) were inferred using the Algorithm for the Reconstruction of Accurate Cellular Networks (ARACNE) with default parameters. The regulons of a potential MR were then divided into positive (upregulated) and negative (downregulated) groups based on Spearman’s correlation coefficients between the expression level of the MR and each gene in its regulon. To test EMT high tumors vs EMT low tumors, a differentially expressed gene (DEG) test was performed, and EMT scores were calculated using the single sample gene set enrichment analysis (ssGSEA) method from the HALLMARK EMT gene set to obtain a ranked differential gene signature (by T statistics). The ranked gene list was used as input for the GSEA method from the R “gage” package to evaluate the enrichment of the regulons, and their associated MRs based on the FDR-adjusted *P* values. Only MRs with significantly enriched associated regulons (cut-off < 0.05) were considered for further experiment validation.

### Cell culture and transfection

The human ovarian cancer cell line HEYA8, OVSAHO, Kuramochi, and A2780 were cultured in RPMI 1640 medium (Sigma-Aldrich), while CaOV3, OVCAR3, and OVCA429 were grown in Medium 199:MCDB105 (Sigma-Aldrich) containing 5% fetal bovine serum (Hyclone) and 1% streptomycin and penicillin (Invitrogen). Cells were cultured in a humidified incubator with 5% CO2 at 37°C. Transient transfection of plasmid DNAs (pEGFPN1 and pEGFPN1-DNM1 wild type) and short interfering RNA were performed using lipofectamine 2000 reagent (Invitrogen) for 48 h according to the manufacturer’s instructions. The plasmid DNAs were purchased from Addgene, and the siRNAs were obtained from Dharmacon. Short hairpin RNA (shRNA) stably expressing cells were generated using lentiviral plasmids carrying DNM1 shRNA (Sigma-Aldrich), followed by selection with 1 μg/ml puromycin (Calbiochem).

### Western blotting

The extracted proteins were separated by sodium dodecyl sulfate-polyacrylamide gel electrophoresis and transferred to the nitrocellulose membranes. The membranes were then blocked with 5% nonfat dry milk and incubated with primary antibodies. Corresponding HRP-conjugated secondary antibodies (Bio-rad) were incubated with the membranes at room temperature for 1 h. After washing 3 times, the membranes were detected using WESTERN LIGHTENING^TM^ Plus-ECL (Perkin Elmer). Antibodies against DNM1, N-cadherin, E-cadherin, vimentin, clathrin, and caveolin-1 were obtained from Cell Signaling Technology. The Dynamin 2 antibody was from Invitrogen, the Dynamin 3 antibody was from Abcam, and the GAPDH antibody was from Immunoway.

### Real time polymerase chain reaction (qPCR)

Total RNA was isolated using Trizol reagent (Invitrogen), followed by reverse-transcription to cDNA using M-MLV reverse transcriptase (Invitrogen) according to the manufacturer’s instructions. qPCR was performed using AceQ qPCR SYBR Green Master Mix (High ROX Premixed) Kit (Vazyme). Relative quantification was calculated by normalizing to the expression level of GAPDH using 2^-ΔΔCt^ method.

### Transwell migration assays

Cells were seeded into 24-well transwell inserts with serum-free medium and allowed to translocate toward the complete medium for 16 h. The migrated cells on the lower surface were fixed with ice-cold methanol and stained with 0.5% crystal violet. For the invasion assay, the inserts were pre-coated with Matrigel (BD Biosciences) and the same procedures were followed. Pictures of 5 random fields from each well were obtained using a microscope at × 10 magnification.

### Surface biotinylation for endocytosis and recycling assays

Cells grown in 6-well plates were treated with 100 ug/ml leupeptin (Roche), 0.5 mM EGTA (Sigma), and 20 ug/ml cycloheximide (Calbiochem) for 30 min at 37°C to block lysosome activity and protein synthesis. Surface proteins were prelabeled with 0.5 mg/ml membrane-impermeable Sulfo-NHS-SS-Biotin (Thermo Scientific) on ice for 15 min. Unbound biotin was quenched with 50 mM NH_4_Cl for 5 min at 4°C twice. Biotinylated cells were rinsed with fresh medium twice and then incubated at 37 °C for 30 min. The cells were quickly chilled to 4°C, and the surface biotin was removed by incubating with 50 mM glutathione solution (75 mM NaCl, 75 mM NaOH, 50 mM reduced glutathione) twice for 15 min on ice. Free glutathione was quenched by 5 mg/ml iodoacetamide twice for 15 min on ice, followed by washing with PBS three times. The cells were then lysed by lysis buffer containing protease inhibitors. Biotinylated proteins were isolated by incubating with NeutrAvidin UltraLink Resin (Thermo Scientific) overnight at 4 °C. The proteins were then separated and analyzed by Western blot. For recycling assays, cells were biotinylated and incubated at 37°C for 30 min and the remaining surface biotinylated proteins were removed by two rounds of glutathione treatment. Free glutathione was quenched by two rounds of iodoacetamide. The cells were rinsed with fresh medium and incubated at 37°C for 15 min, followed by two rounds of glutathione treatment and iodoacetamide treatments.

### Immunofluorescence staining

For the colocalization assay, cells grown in 6-well plates were treated with 0.5 mM EGTA and 20 ug/ml cycloheximide for 30 min at 37°C to disrupt N-cadherin homophilic interactions and block protein synthesis. The cells were then incubated with antibody against the extracellular domain of N-cadherin (Invitrogen) for 1 h at 4°C. After the antibody incubation, cells were washed twice with fresh medium to remove unbound antibody, followed by incubation at 37°C for 15 min. After fixation with 4% paraformaldehyde, cells were blocked with 5% bovine serum albumin containing 0.5% Triton X-100 for 30 min prior to staining with Rab11 primary antibody overnight at 4°C. The cells were washed twice with PBS and further incubated with Cy3-conjuated anti-rabbit secondary antibodies and Cy5-conjuated anti-mouse secondary antibodies (Bio-rad) for 1 h. Cells were stained with Hoechst33342 to visualize the nuclei. The stained cells were mounted on glass slides with Vectashield mounting medium (Vecta Laboratories) and imaged with Zeiss ELYRA S1 super resolution microscope. For cell polarity analysis, cells were seeded on glass cover slips and cultured to full confluence in the presence of silicone inserts to create wound (Ibidi). The cells were further cultured for 6 h after insert removal, followed by fixation and staining with anti-N-cadherin, anti-GM130 and Hoechst 33342, and imaged by a Carl Zeiss LSM 900 confocal microscope.

### Sequencing

Total RNA was extracted from HM and NM cells with Trizol Reagent (Invitrogen). RNA-seq was performed at the Centre for PanorOmic Sciences (CPOS), the University of Hong Kong. For ATAC-seq, the Omni-ATAC-seq method was used with minor modifications (Corces, et al., 2017). Briefly, 50,000 viable cells were centrifuged at 500 g at 4°C for 5 min. The cell pellet was resuspended with 50 μL cold ATAC-Resuspension Buffer (RSB) (5 M NaCl, 1 M MgCl2, 10 mM Tris HCl pH 7.4 and protease inhibitor cocktail) containing 0.1% Igepal CA-630, 0.1% digitonin. After incubation on ice for 3 min, the sample was washed with 1 mL cold ATAC-RSB containing 0.1% tween-20, followed by centrifugation at 500 g for 10 min at 4°C. Tagmentation was performed by adding 50 μL transposase mix (16.9 μL PBS, 2.5 μL Tn5, 25 μL 2X TD buffer, 0.1 μL 5% digitonin, 0.5 μL 10% Tween-20 and 5 μL diH2O) and incubating in a thermocycler (1000 rpm) at 37°C for 30 min. Tagmented DNA was purified using the MinElute Kit. Library amplification was performed using indexed primers with 50 μL Kapa Hi Fi Hot Start PCR reaction for the first 5 cycles. 1 μL of amplified product was used to perform a library quantification with the KAPA library quantification kit to determine the optimum number of final amplification cycles. AMPure XP beads were used to select for fragments of 150 bp to 800 bp and the library size was determined by Fragment Analyzer. The obtained library was then sent to Novogene for library quality check, quantification, and sequencing.

### Mice experiments

All mouse experiments were conducted in accordance with the guidelines of the University of Hong Kong Animal Care and Use Committee. Luciferase-expressing HM cells treated with NS or DNM1 shRNA (1×10^6^) were injected intraperitoneally into 4-week-old female NOD/SCID mice (n=3/group). The mice were imaged twice a week for 22 days using the Xenogen IVIS system. The tumors on the omentum were harvested and metastatic nodules in the peritoneal cavity were counted. Formalin was used for tissue specimen fixation, and the expression of N-cadherin and vimentin were stained using corresponding antibodies.

### Immunohistochemistry (IHC)

FFPE sections were subjected to deparaffinization with xylene for 5 min three times and rehydration with graded ethanol (absolute ethanol, 95%, 70% and 50%, twice each). Antigen retrieval was performed using citrate buffer. After blocking, the slides were incubated with primary antibodies against PAX8, N-cadherin, and vimentin (Cell Signaling Technology), and negative control at 4°C overnight. The slides were further processed using rabbit specific HRP/DAB (ABC) detection IHC kit according to the manufacturer’s protocol (Abcam).

### siRNA dendriplexes formation

AmDM (MW=3838 g/mol, 16 amine end groups) was dissolved in distilled H_2_O and stocked at 500 μM. To prepare the siRNA dendriplexes, the dendrimer and siRNA were first diluted separately in OPTI-MEM (Invitrogen) and incubated at room temperature for 5 min. Dendrimer and siRNA (20 nM) were then mixed and incubated at room temperature for 20 min before treatment.

### Statistics

Data were shown as mean ± SD and statistical analyses were conducted using student’s t-test by GraphPad (San Diego, CA) for comparison between two groups. For comparisons involving three or more groups, one-way ANOVA was used for statistical analysis. *P* < 0.05 was considered statistically significant.

## ACKNOWLEDGMENTS

We thank Prof Danny C. Y. Leung (Hong Kong University of Science and Technology) and Dr Sophia S. N. Lam for providing assistance in the analysis of the ATAC-seq data. This work was supported by the Senior Research Fellow Scheme SRFS2223-7S05 to A. S. T. Wong, and from “Laboratory for Synthetic Chemistry and Chemical Biology” under the Health@InnoHK Program launched by Innovation and Technology Commission, HKSAR. This work was supported by the Hong Kong Scholars Program [XJ2019051].

## AUTHOR CONTRIBUTIONS

A.S.T.W., Y. C. and S.KY.T. designed the research and prepared the manuscript. Y. C. and Z. G. performed research experiments and analyzed data. Y. T. and J.Z. performed the bioinformatics analysis. P.P.C.I. provided clinical samples. L.P. provided dendrimers. A.S.T.W. supervised the study.

