## Supplementary Figures for "Dynamin 1-mediated endocytic recycling of glycosylated N-cadherin sustains the plastic mesenchymal state to promote ovarian cancer metastasis"

### Supplementary Figure 1

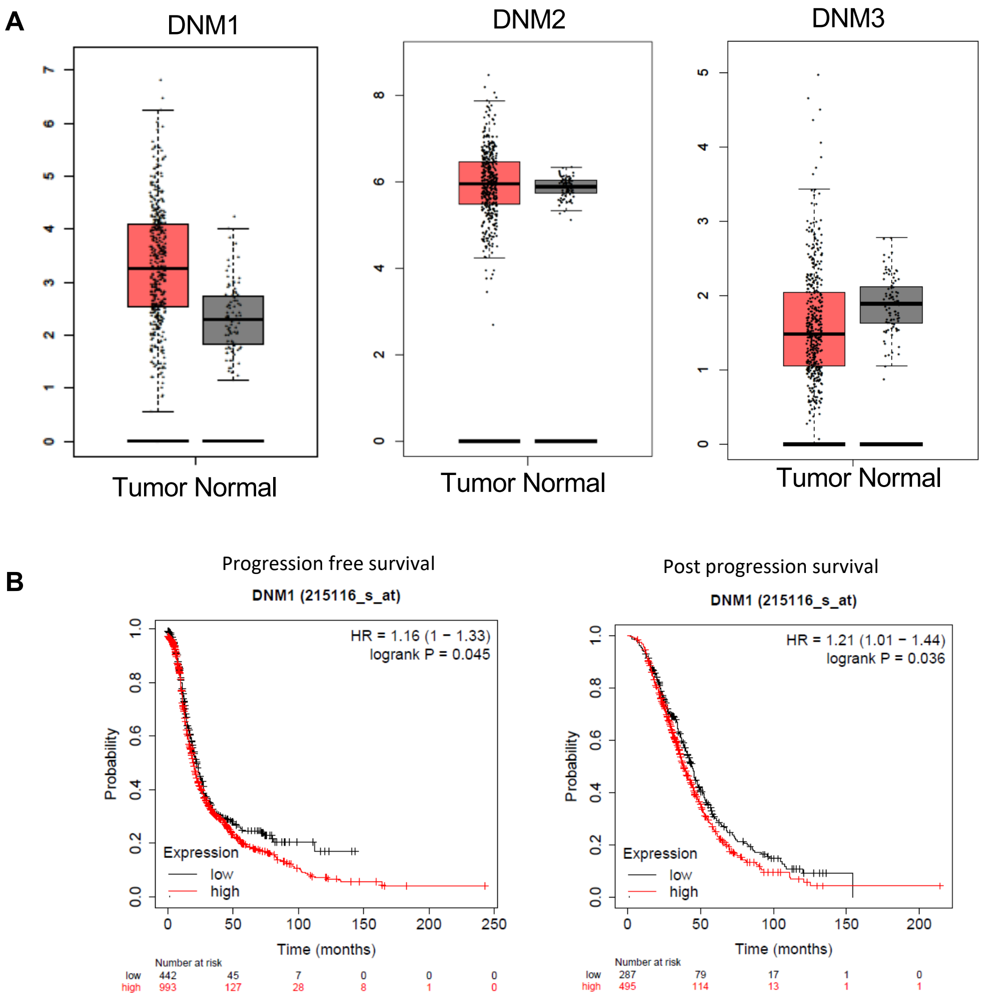


**Supplementary Figure 1**. **(A)** The mRNA expression level of DNM1, DNM2 and DNM3 in ovarian cancer (n=426) and normal tissues (n=88) were shown. **(B)** Progression free survival and post progression survival of patients stratified by DNM1 expression level in ovarian cancer were shown.

### Supplementary Figure 2

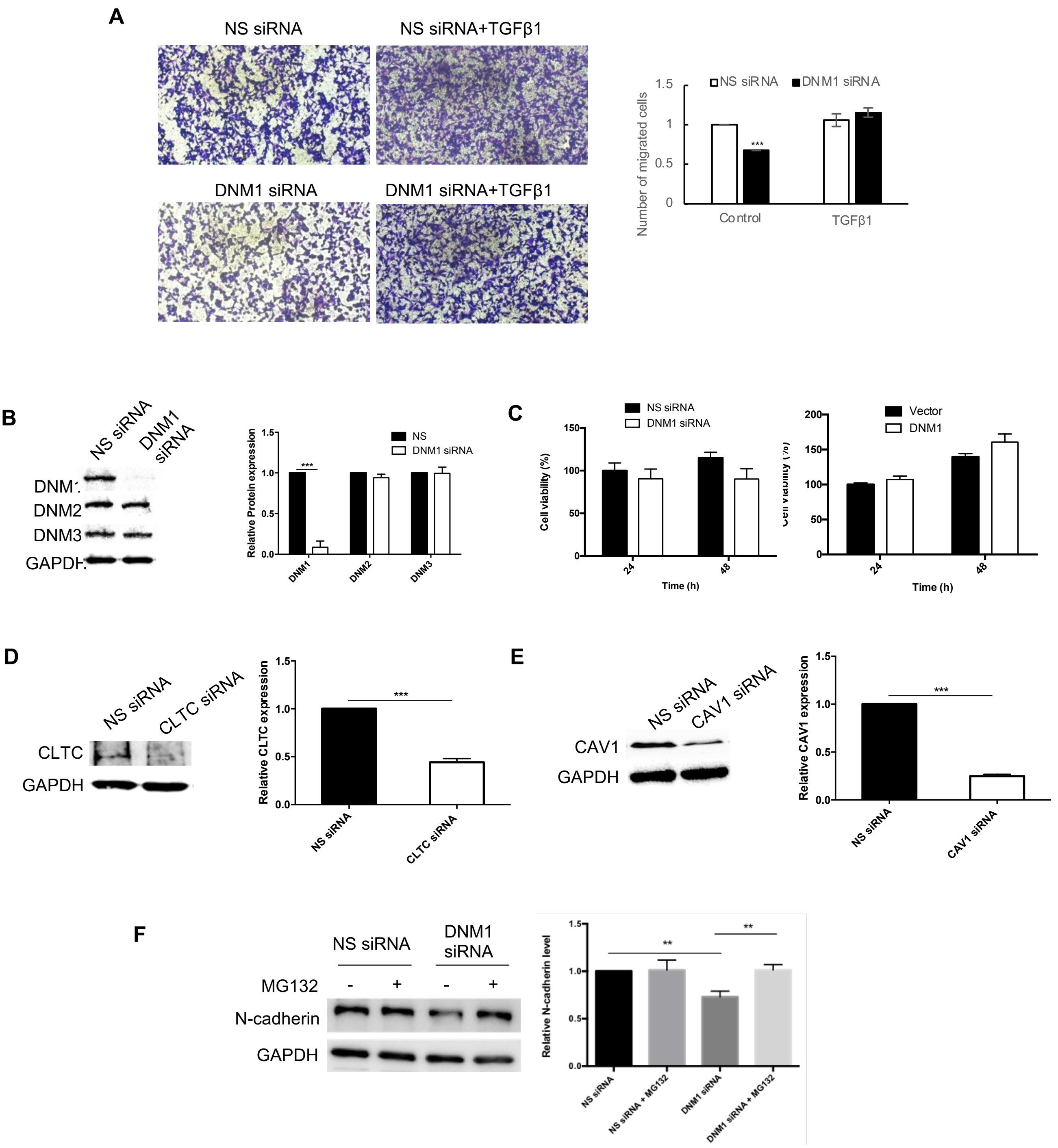


**Supplementary Figure 2**. **(A)** HM cells transfected with NS or DNM1 siRNA were treated with or without TGF- β as indicated. **(B)** DNM1, DNM2 and DNM3 expression were tested in HM cells transfected with NS or DNM1 siRNA. **(C)** MTT cell viability assays were performed for HM cells treated with NS or DNM1 siRNA and NM cells transfected with control or DNM1 overexpression vector. **(D)** HM transfected with NS or clathrin (CLTC) siRNA was tested for clathrin expression by Western blot. **(E)** HM transfected with NS or caveolin-1 (CAV1) siRNA was tested for caveolin expression by Western blot. **(F)** HM transfected with DNM1 siRNA were exposed to MG132 (50 uM) for 4 h and expression level of N-cadherin was detected by Western blot. Band intensities were quantified by imageJ and GAPDH served as a loading control. Representative blots of three independent experiments were shown. *, $P < 0.05$. **, $P < 0.01$ and ***, $P < 0.005$.

### Supplementary Figure 3

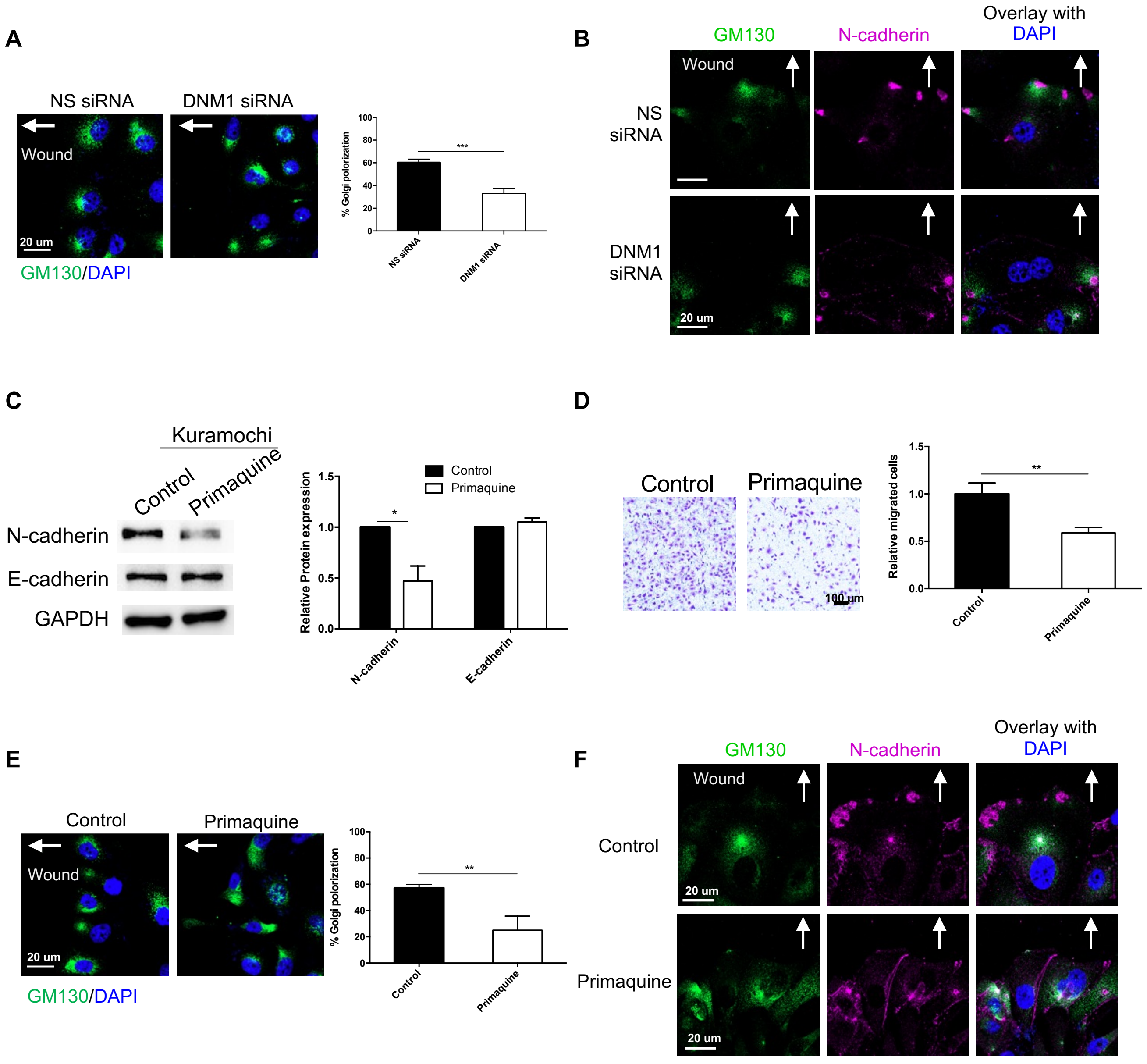


**Supplementary Figure 3**. **(A)** Cell-free gaps were created in Kuramochi monolayers after transfection with NS or DNM1 siRNA. Golgi orientation was visualized by staining the Golgi marker GM130 (green). The number of cells with correct polarity were counted. **(B)** Cells were co-stained with GM130 and N-cadherin (magenta), followed by observation by confocal imaging. Kuramochi cells were treated with vechicle control or primaquine (100 µM). Treated cells were analysed by the following assays. **(C)** N-cadherin expression was tested by Western blot. (B) Biotin recycling assay was performed, followed by Western blot for N-cadherin. **(D)** Migration assays was performed. Migrated cells were stained with crystal violet and counted in five random fields of view.**(E)** Cell-free gaps were created in the cell monolayer. Golgi orientation was visualized by staining the Golgi marker GM130 (green). The number of cells with correct polarity were counted. **(F)** Treated cells were co-stained with GM130 and N-cadherin (magenta), followed by confocal imaging. Nuclei were counterstained by DAPI (blue). Arrows indicated the expected Golgi direction towards the wound. Representative data of three independent experiments were shown. *, $P < 0.05$. **, $P < 0.01$ and ***, $P < 0.005$.

### Supplementary Figure 4

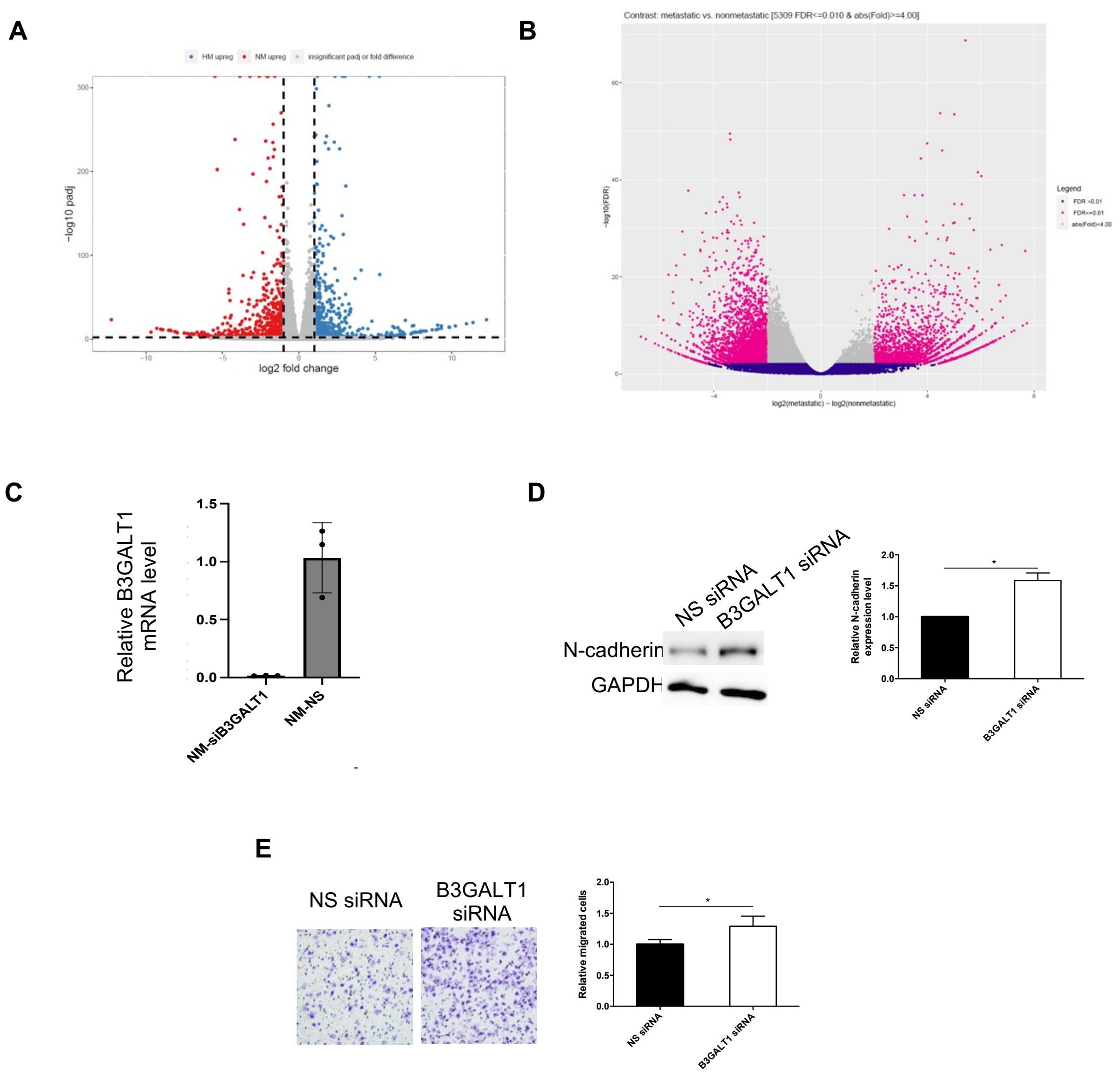


**Supplementary Figure 4**. Volcano plots of **(A)** differentially expressed genes (by RNA-seq) and **(B)** differentially accessible peaks (by ATAC-seq) between NM and HM were shown. **(C)** HM cells transfected with NS or B3GALT1 siRNA were tested for B3GALT1 expression by RT-PCR. **(D)** N-cadherin expression was tested by Western blot in Kuramochi transfected with NS or B3GALT1 siRNA. GAPDH was used as a loading control and band intensities were quantified by ImageJ. **(E)** Migration assays were performed in Kuramochi transfected with NS or B3GALT1 siRNA. Migrated cells were stained with crystal violet and counted in five random fields of view. Representative data of three independent experiments were shown. *, $P < 0.05$. **, $P < 0.01$ and ***, $P < 0.005$.
